# Loss of the Pioneer Factor FOXA1 Results in Genome-wide Epigenetic Reprogramming and activation of Interferon-Response Genes including *CD274*/PD-L1

**DOI:** 10.1101/2020.09.11.294041

**Authors:** Wenhuo Hu, Hironobu Yamashita, Jenna Craig, Vonn Walter, Joshua I. Warrick, Hikmat Al-Ahmadie, David J. DeGraff

**Affiliations:** Human Oncology and Pathogenesis Program, Memorial Sloan Kettering Cancer Center, New York, NY 10065; Department of Pathology and Laboratory Medicine, Penn State College of Medicine, Hershey, PA 17033; Department of Surgery, Division of Urology, Penn State College of Medicine, Hershey, PA 17033; Department of Public Health Sciences, Penn State College of Medicine, Hershey PA 17033; Department of Pathology, Memorial Sloan Kettering Cancer Center, New York, NY 10065

## Abstract

Forkhead Box A1 (FOXA1) is a pioneer transcription factor critical in epigenetic regulation of chromatin and cell fate determination. Reduced FOXA1 expression is an independent predictor of poor overall survival in bladder cancer patients. However, the impact of *FOXA1* loss on chromatin epigenetics in bladder cancer is unknown. Therefore, we determined the impact of *FOXA1* knock out (KO) on epigenetic modification of chromatin and associated gene expression. We identified 8,230 differentially expressed genes following *FOXA1* KO. Surprisingly, Gene Set Enrichment Analysis (GSEA) identified IFNɑ/ɣ gene expression signatures as enriched following *FOXA1* KO. *FOXA1* KO induced both increased and decreased numbers of histone 3 lysine 27 acetylation (H3K27ac) sites throughout the genome. As expected, the majority of differences in H3K27ac across genomic areas in *FOXA1* KO cells is mapped to intergenic and intronic regions where enhancers reside. In addition, a subset of differential H3K27ac levels were also mapped to proximal promoters and within gene bodies. Integrated analysis of RNA/ChIP-seq data shows changes in gene expression that are mirrored by differences in H3K27ac. Motif analysis of DNA sequence enriched for H3K27ac identified significant increases in transcription factor binding motifs including the interferon sensitive response element (ISRE) and interferon response factors such as IRF1. Moreover, we identified increased H3K27ac of regulatory elements as being associated with several upregulated interferon sensitive genes (ISGs) in *FOXA1* KO cells, including *CD274*/PD-L1. Western blotting and Q-RT-PCR confirmed upregulation of *CD274*/PD-L1 following *FOXA1* KO. Analysis of TCGA data confirmed an inverse relationship between *FOXA1* and *CD274* in bladder cancer, as well as in other cancers. In summary, we provide evidence of widespread epigenetic reprogramming after *FOXA1* KO in bladder cancer cells. Additionally, we provide evidence that *FOXA1* KO-induced epigenetic changes contribute to activation of a global interferon-dominant expression signature, including the immune checkpoint target *CD274*/PD-L1 in a cancer cell-intrinsic manner.

## Introduction

We identified loss of Forkhead Box A1 (FOXA1) expression as a common event in the development of bladder cancers with squamous differentiation^1^, and subsequently showed that FOXA1 is an independent predictor of poor overall survival in this common malignancy^2^. Additional *in vitro* studies showed that FOXA1 overexpression cooperates with overexpression of GATA Binding Protein 3 (GATA3) and pharmacologic activation of Peroxisome Proliferator Activated Receptor Gamma (PPARɣ) to repress a basal-squamous signature *in vitro*^3^. Moreover, we went on to show that *in vivo* ablation of *FOXA1* results in the development of squamous differentiation and cooperates with other genetic events resulting in the development of bladder cancer^4^. While our observations have identified an important role for this pioneer factor in bladder cancer, the mechanism by which FOXA1 impacts gene expression in bladder cancer remains unknown.

In addition to performing a role in organogenesis and tissue-specific gene expression, pioneer transcription factors such as FOXA1 directly bind to areas of condensed chromatin prior to recruitment of other coregulatory factors^5, 6^. Indeed, FOXA1-chromatin binding plays an important biologic role for organogenesis and tissue-specific gene expression^7^. Because of the structural similarity of the FOXA1 DNA binding domain to linker histone^8^, FOXA1 can displace linker histone, resulting in increases in local DNA accessibility to enable binding of additional *trans* acting factors^9^. In addition, FOXA1 binding to chromatin can result in the “passive” reduction in the number of additional binding events required for transcriptional activation^9^. Moreover, bookmarking of specific genomic loci by FOXA1 during mitosis enables rapid expression of genes following mitotic exit, thus preserving cell identify^10^. Interestingly, active and passive pioneering, as well as bookmarking by FOXA1 are accompanied by epigenetic changes in chromatin modification, which are in turn associated with DNA accessibility by trans-acting factors, as well as gene expression changes.

In addition to a role in organogenesis and tissue-specific gene expression, ample evidence indicates a role for FOXA1-mediated regulation of chromatin accessibility in disease pathogenesis. For example, alterations in FOXA1 expression or mutational status have been associated with chromatin alterations in hormonally-regulated cancers of the breast and prostate^11–14^, as well as other malignancies including liver^15^ and pancreatic cancer^16^. Therefore, we hypothesized that decreased expression of FOXA1 in urothelial cancer cells has an important, and disease-relevant impact on epigenetic post-translational modification of chromatin.

## Results

### FOXA1 is a cancer cell-intrinsic repressor of the IFNα and IFNɣ transcriptional signature including *CD274***/PD-L1 expression**

To gain insight into alterations in gene expression associated with *FOXA1* inactivation, we genetically ablated *FOXA1* in luminal UMUC1 bladder cancer cells, resulting in the creation of two separate knockout (KO) clones. Principal component analysis performed on RNA seq data identified a clear separation between parental and *FOXA1* KO cells (Figure 1A). Differential expression analysis identified 8,230 differentially expressed genes following *FOXA1* KO (Figure 1B; 4,185 increased, 4,045 decreased; FDR q<0.05; two separate KO clones, average of differentially expressed genes in both clones). Interrogation of upregulated genes from individual UMUC1 *FOXA1* KO clones via Gene Set Enrichment Analysis (GSEA) analysis strongly suggested a role for FOXA1 in repression of interferon target genes (Figure 1C). For example, GSEA identified both “interferon alpha (IFNɑ) response” (*FOXA1* KO clone 1; GSEA pre-rank analysis FDR q = 0.005, NES = 1.79; *FOXA1* KO clone 2, GSEA pre-rank analysis FDR q < 0.001, NES = 2.54) and “interferon gamma (IFNɣ) response” (*FOXA1* KO clone 1; GSEA pre-rank analysis FDR q = 0.04, NES = 1.42; *FOXA1* KO clone 2, GSEA pre-rank analysis FDR q < 0.001, NES = 2.02) as being enriched following *FOXA1* ablation (Figure 1C). Amongst genes expressed at significantly lower levels following *FOXA1* KO, enriched gene sets included “MYC targets V1”, “E2F targets” and “androgen response” (data not shown). This latter finding is noteworthy because of the fact that FOXA1 is known to interact with the androgen receptor to dictate the androgen responsive cistrome^17–19^. We next focused the expression analysis on genes relevant to interferon signaling, and found *FOXA1* KO cells demonstrated significantly increased expression of *CD274*, the gene encoding the immune checkpoint target Programmed Death-Ligand 1 (Figure 1D; q<0.05 for both *FOXA1* KO clone 1 and 2). This is in keeping with the fact that FOXA1 has been previously identified as a direct, positive regulator of *CD274/*PD-L1 in a subset of T regulatory cells^20^. In addition, *FOXA1* KO resulted in significantly increased expression of several other interferon-sensitive genes (ISGs), including *IFI35*, *IRF9*, *IFIT2* and *STAT2* among others (Figure 1D; q<0.05 for both *FOXA1* KO clone 1 and 2).

**Figure 1:**
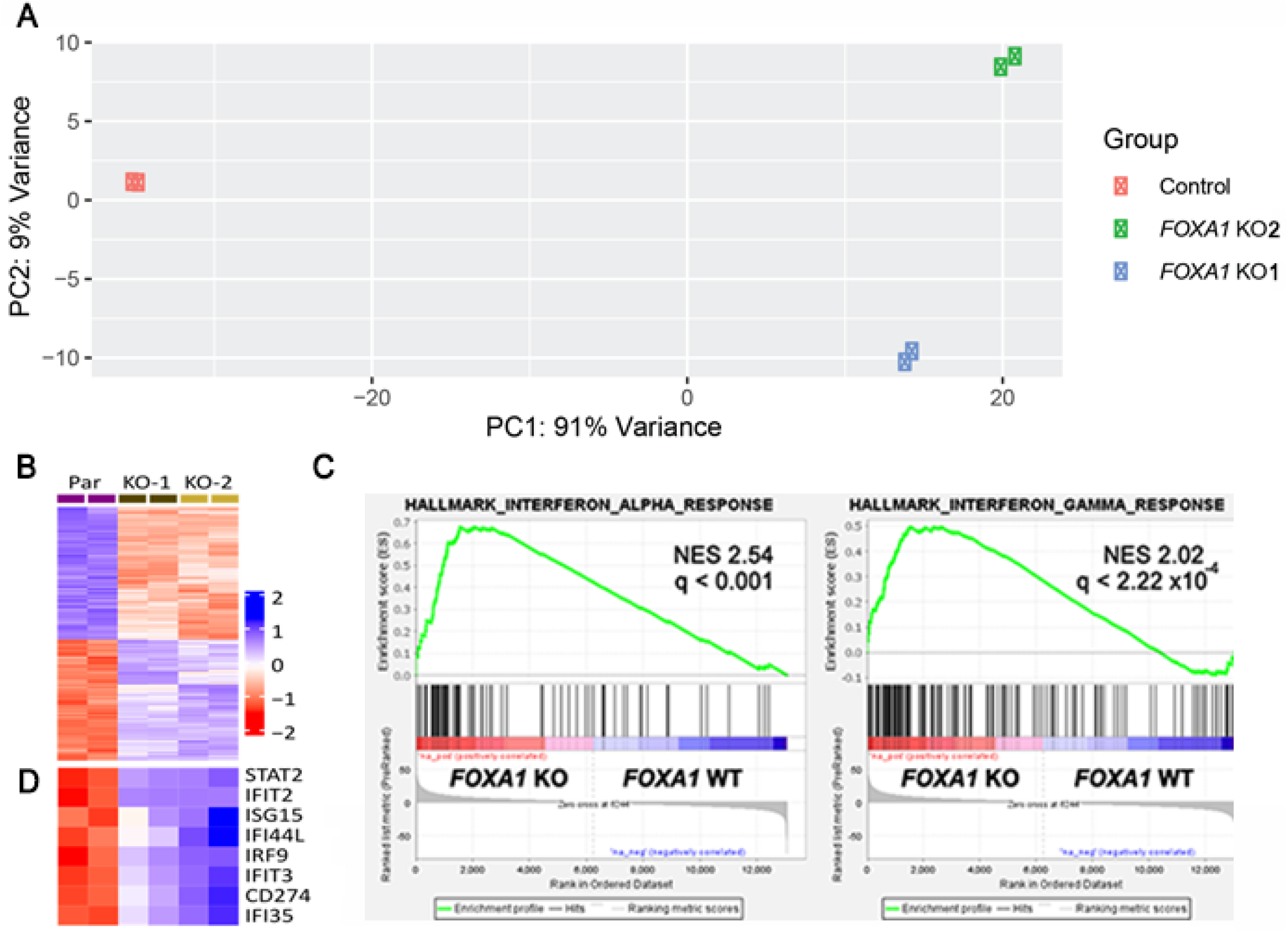
*FOXA1* knockout in UMUC1 bladder cancer cells activates a global IFNα/ɣ gene expression program associated with response to immune checkpoint treatment. (**A**) Principal components analysis plot illustrates differences in gene expression patterns observed in two *FOXA1* knockout UMUC1 and control UMUC1 bladder cancer cells. Technical replicates cluster together, illustrating highly concordant gene expression patterns between the technical replicates. In contrast, both *FOXA1* knockout UMUC1 clones are markedly different from the parental UMUC1 clones (primarily PC1), and also different from each other (primarily PC2). (**B**) Heat map showing 1,914 differentially expressed genes (FDR q<0.05) following *FOXA1* KO in UMUC1. Two replicates are shown for each sample, with two biologic replicates for UMUC1 *FOXA1* KO. (**C**) Gene Set Enrichment Analysis (GSEA) identified enrichment for genes included in the Hallmark (**A**) IFNα and IFNɣ response in *FOXA1* KO UMUC1 cells (clone 2 shown; see text for NES and q values for clone 1). (**D**) Differentially expressed interferon-sensitive genes (ISGs) are highlighted.

Based on these results, we further investigated the impact of altered FOXA1 levels on *CD274*/PD-L1 expression. Q-RT-PCR (Figure 2A and B; Student’s t-test; p < 0.05) and western blotting (Figure 2C, D and E; Student’s t-test; p < 0.05) of parental UMUC1 and UMUC1 *FOXA1* KO cells confirmed significantly increased expression of *CD274*/PD-L1 following *FOXA1* KO. The ability of FOXA1 to regulate *CD274*/PD-L1 was additionally supported by FOXA1 gain of function studies. Specifically, we identified significant decreases in *CD274*/PD-L1 expression following FOXA1 overexpression in UMUC3 bladder cancer cells (Figure. 2F thru J; Student’s t-test; p < 0.05). Analysis of publicly available data from TCGA confirmed an inverse relationship between *FOXA1* and *CD274* expression in a cohort of bladder cancer patients (Figure 2K; Spearman correlation; r= −0.45; p < 0.001). In addition to bladder cancer, analysis of pan-cancer TCGA data identified a statistically significant associations between *FOXA1* and *CD274* expression in a number of malignancies (Figure 2L). In summary, these findings identify FOXA1 as a cancer cell-intrinsic repressor of *CD274*/PD-L1 in bladder cancer, as well as potentially in other malignancies.

**Figure 2:**
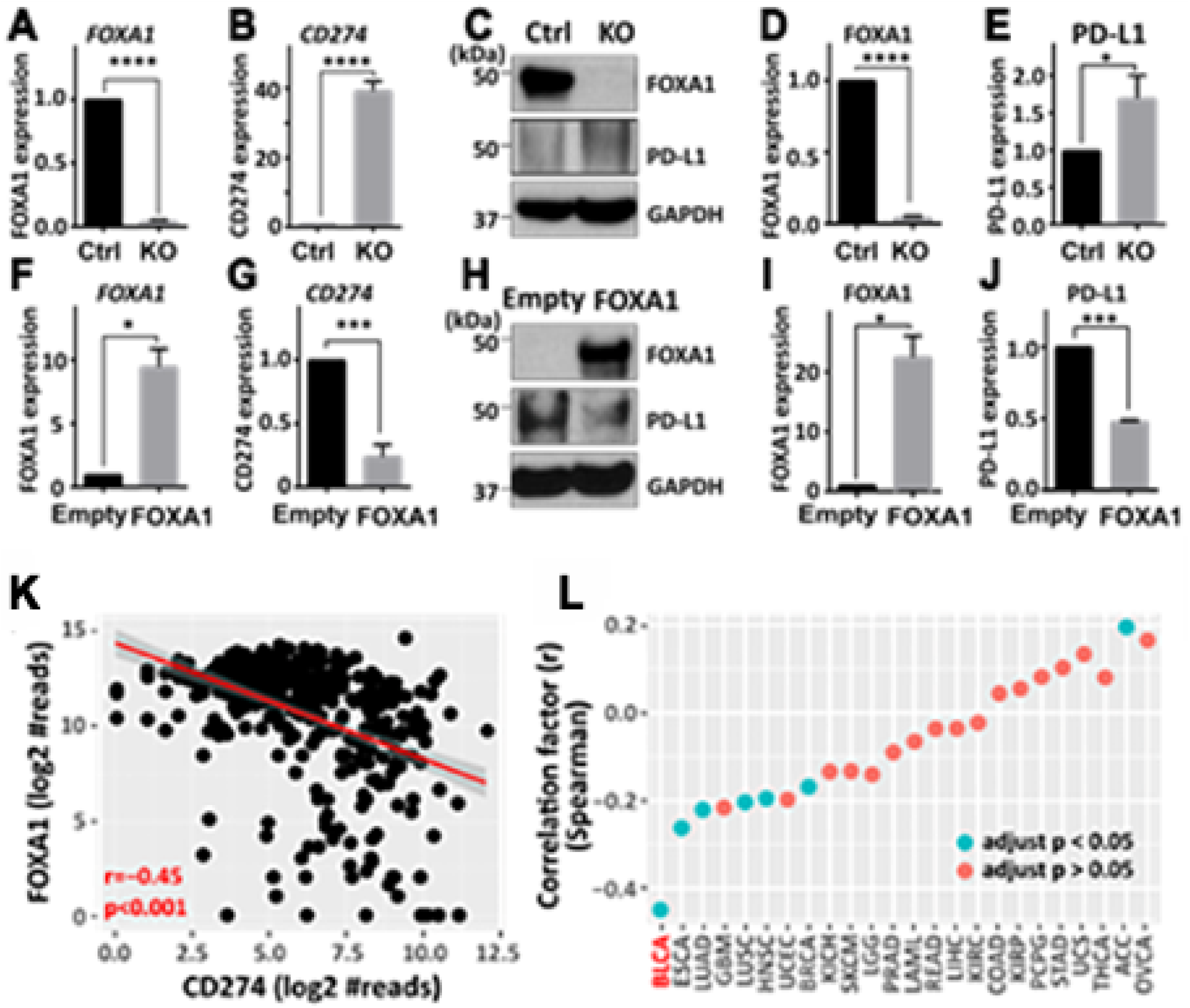
*CD274*/PD-L1 is a FOXA1-repressed immune checkpoint target. Q-RT-PCR (**A** and **B**), western blotting (**C**) and densitometry (**D** and **E**) confirms decreased FOXA1 and increased *CD274*/PD-L1 expression in UMUC1 *FOXA1* KO cells. Q-RT-PCR (**F** and **G**), western blotting (**H**) and densitometry (**I** and **J**) confirms increased FOXA1 and decreased *CD274* /PD-L1 expression in UMUC3 *FOXA1* overexpressing cells. (**K**) Inverse correlation between *FOXA1* and *CD274* (encoding PD-L1 expression) in the TCGA bladder data set (r=−.45; p<0.001; Spearman correlation test). (**L**) Spearman correlation of *FOXA1* and *CD274* across cancers identifies a significant correlation between these genes in a number of malignancies.

### Activation of the interferon-dominant immune signature and *CD274*/PD-L1 is a result of genome-wide epigenetic reprogramming following *FOXA1* ablation

The ability of FOXA1 to regulate chromatin accessibility at enhancer and promoter elements via displacement of linker histone is a hallmark characteristic of this and other pioneer factors^5, 9^. Genome-wide changes in one mark of active enhancers and promoters, Histone 3 lysine 27 acetylation (H3K27ac), has been reported following changes in FOXA1 expression in a number of malignancies^16, 21^. While the impact of *FOXA1* silencing on the epigenomic landscape in bladder cancer has not been reported, we hypothesized that increased expression of *CD274* and other ISGs may be a result of changes in H3K27ac of associated genes following *FOXA1* loss. To directly test this hypothesis, we subjected parental and UMUC1 *FOXA1* KO (clone 2; two technical replicates) to chromatin immunoprecipitation coupled to high throughput sequencing (ChIP-seq) for H3K27ac, and integrated these results with our RNA-seq data following *FOXA1* KO. There were 14,977 locus with H3K27ac enrichment in either parental UM-UC-1 cells or its FOXA1 KO cells (FDR < 0.05). Consistent with the accepted role of FOXA1 in the control of enhancer activity, the majority of differences in H3K27ac areas following *FOXA1* KO were mapped to intergenic (n=6,250 peaks; 42% of total peaks) and intronic (n= 6,490 peaks; 43%) genomic regions (Figure 3A). In addition, differential H3K27ac levels following *FOXA1* KO were also mapped to proximal promoters (n= 1,306 peaks; 9%) as well as within gene bodies (931 peaks; 6%).

**Figure 3:**
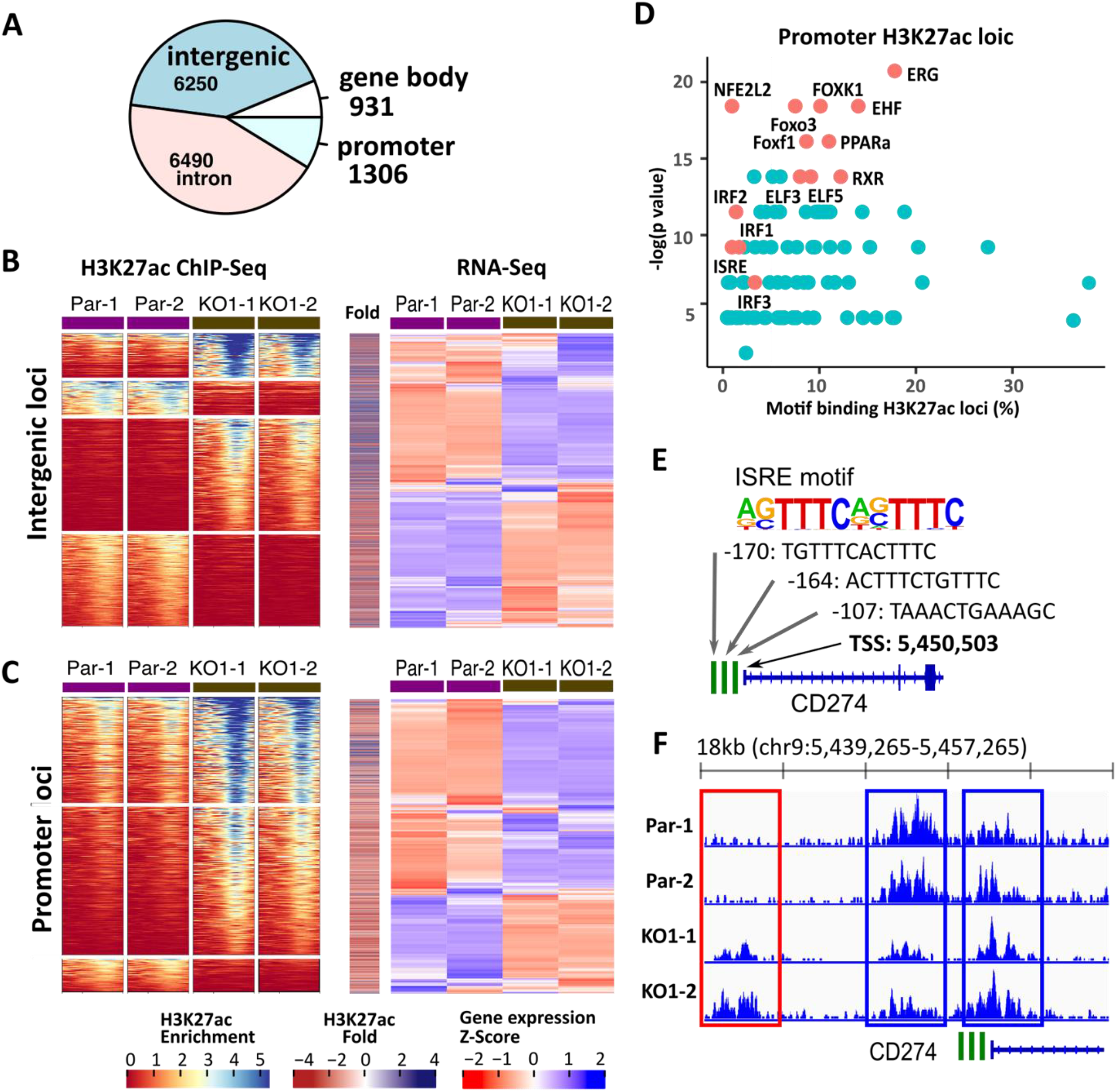
Genetic knockout of *FOXA1* results in widespread enhancer and promoter reprogramming in human cells. (**A**) Genomic region annotation of H3K27ac modification locus. (**B** and **C**) H3K27ac cluster analysis (left panels) and expression heatmaps (right panels) showing relationship between H3K27ac ChIP-seq and expression of 1,724 associated genes (RNA-seq) at (**B**) intergenic (including enhancers) and (**C**) promoter levels. These data show decreased H3K27ac is associated with reduced gene expression following *FOXA1* KO, while increased H3K27ac is associated with increased gene expression following *FOXA1* KO. (**D**) Motif analysis identifying enrichment of binding sites for transcription factors activated by IFNα/ɣ signaling at areas of increased H3K27ac following *FOXA1* KO. (**E**) Illustration identifying the location of several interferon-sensitive response element (ISRE) motifs in the areas of increased acetylation in the *CD274* promoter. (**F**) Increased acetylation of *CD274* regulatory elements including an upstream enhancer (red box) and the proximal promoter region (blue boxes) following *FOXA1* KO.

Within the 6,250 H3K27ac peaks enriched in intergenic region, we identified 4 distinct clusters (Figure 3B). Clusters 1 (n=943 peaks) and 3 (n=2499 peaks) exhibited significant increases in H3K27ac levels, while clusters 2 (n=883 peaks) and 4 (n=1925 peaks) exhibited significant decreases in H3K27ac marks at intergenic loci. Confirming an important role for FOXA1 in the regulation of enhancer activity, integrated RNA-seq and ChIP-seq results identified significant changes in gene expression mirrored differences in H3K27ac (Figure 3B). While intergenic regions showed both significant numbers of decreased and increased H3K27ac marks following *FOXA1* KO, we identified widespread increases in H3K27ac marks at proximal promoters (Figure 3C). Specifically, 3 separate clusters were identified, with clusters 1 (n=484 peaks; 37% of promoter peaks) and 2 (n=664 peaks; 51% of promoter peaks) exhibiting increases in H3K27ac marks, while cluster 3 exhibited a reduction in H3K27ac marks (n=158 peaks; 12% of promoter peaks). In keeping with increased levels of H3K27ac at promoters, expression of greater than half of genes in clusters 1 and 2 were upregulated following RNA-seq analysis (Figure 3C). These results suggest an important, yet perhaps underappreciated role for FOXA1 in the control of proximal promoter activity. As H3K27ac marks areas of open chromatin associated with active regulatory elements^22, 23^, we performed motif analysis in an effort to determine what transcription factors potentially bind to newly accessible areas following *FOXA1* KO. In support of a role for *FOXA1* KO in the global activation of ISGs, enriched transcription factor binding motifs included the interferon sensitive response element (ISRE; q<0.05), as well as motifs for interferon response factors (IRF1, IRF2 and IRF3; all q<0.05) (Figure 3D). Focusing on the *CD274* proximal promoter, we identified 3 separate ISRE elements in this region (Figure 3E). As predicted by our motif analysis, these three ISRE elements overlapped with areas of H3K27ac following in both parental and *FOXA1* KO UMUC1 cells, which suggests basal promoter activity (Figure 3F). In addition, we identified an additional regulatory element upstream of *CD274* as having increased H3K27ac following *FOXA1* KO (Figure 3F). In summary, we provide evidence of widespread epigenetic reprogramming following *FOXA1* KO in bladder cancer cells. Additionally, we provide evidence that epigenetic changes contribute to activation of a global interferon-dominant signature, including *CD274*/PD-L1 in a cancer cell-intrinsic manner following *FOXA1* inactivation.

## Discussion

This short report identifies FOXA1 as a repressor of CD274/PD-L1 expression, indicating a tumor cell-intrinsic contribution to PD-L1 expression. Increased H3K27ac is an epigenomic mark of increased activity of *cis* regulatory elements^22, 23^. We found that FOXA1 inactivation *in vitro* drives genome-wide epigenetic reprogramming via alterations in chromatin acetylation, specifically increasing H3K27ac at regulatory elements important for the control of *CD274*/PD-L1 and other ISGs. The identification of FOXA1 as a tumor cell-intrinsic repressor of *CD274*/PD-L1 expression was further supported by gain of function studies. Mutations in *FOXA1* have been associated with activation of IFNɣ-target genes in prostate cancer^12^, and FOXA1 has previously been implicated in the positive control of PD-L1 expression in a subset of immune cells^20^. However, to the best of our knowledge this is the first report of FOXA1 as a tumor cell-intrinsic repressor of PD-L1 in any malignancy. FOXA1 is a recognized pioneer factor controlling chromatin structure^5, 6^, and our findings suggest global epigenetic reprogramming following *FOXA1* loss contributes to the activation of IFN-dominant signatures. This suggestion is supported by motif analysis following H3K27ac ChIP-seq, which demonstrated that areas of acetylated DNA following *FOXA1* loss are enriched for binding sites of a number of interferon-responsive factors. This finding suggests that *FOXA1* inactivation “primes” tumor cells, making them more sensitive to paracrine interferons produced by infiltrating inflammatory cells. Furthermore, the possibility that FOXA1 serves as an important regulator of PD-L1 in other malignancies is supported by our retrospective analysis of TCGA data. More work is required to determine the functional implications increased ISGs and PD-L1 expression for infiltrating immune cells.

## Materials and Methods

### Cell culture and generation of FOXA1 knockout lines

Human bladder cancer cells (UMUC1, UMUC3) were purchased as described previously^3^, and authenticity was confirmed by short tandem repeat (STR) analysis. Cells were cultured in Minimal Essential Medium (UMUC1, UMUC3) supplemented with 10% FBS. To establish UMUC1-FOXA1 knockout (KO), UMUC1 (2×10^5^) cells were transfected with 2.5 mg of HNF-3alpha CRISPR/Cas9 KO plasmid (Santa Cruz, sc-400743) using lipofectamine3000 (Thermo fisher scientific). After 48 h, transfected cells were trypsinized and resuspended in PBS. Three GFP-positive cells were FACs sorted in single well of 96-well plate containing 100 ml of medium. Sorted cells were expanded and sequentially transferred to 24 well, 6 well dishes, and T-75 flasks. Finally, knockout of *FOXA1* in UMUC1-FOXA1KO cells was confirmed by qPCR and western blot analysis.

### ChIP-Seq

Cells were fixed with 1% formaldehyde for 15 min and quenched with 0.125 M glycine. Chromatin was isolated by the addition of lysis buffer, followed by disruption with a Dounce homogenizer. Lysates were sonicated, and the DNA sheared to an average length of 300-500 bp. Genomic DNA (Input) was prepared by treating aliquots of chromatin with RNase, proteinase K and heat for de-crosslinking, followed by ethanol precipitation. Pellets were resuspended, and the resulting DNA was quantified on a NanoDrop spectrophotometer. Extrapolation to the original chromatin volume allowed quantitation of the total chromatin yield. An aliquot of chromatin (30 ug) was precleared with protein A agarose beads (Invitrogen). Genomic DNA regions of interest were isolated using 4 ug of antibody against H3K27Ac (Active Motif, cat# 39133, Lot# 28518012). Complexes were washed, eluted from the beads with SDS buffer, and subjected to RNase and proteinase K treatment. Crosslinks were reversed by incubation overnight at 65 C, and ChIP DNA was purified by phenol-chloroform extraction and ethanol precipitation. Quantitative PCR (QPCR) reactions were carried out in triplicate on specific genomic regions using SYBR Green Supermix (Bio-Rad). The resulting signals were normalized for primer efficiency by carrying out QPCR for each primer pair using Input DNA. Illumina sequencing libraries were prepared from the ChIP and Input DNAs by the standard consecutive enzymatic steps of end-polishing, dA-addition, and adaptor ligation. Steps were performed on an automated system (Apollo 342, Wafergen Biosystems/Takara). After a final PCR amplification step, the resulting DNA libraries were quantified and sequenced on Illumina’s NextSeq 500 (75 nt reads, single end). Reads were aligned to the human genome (hg38) using the Bowtie2 algorithm (default settings). Duplicate reads were removed and only uniquely mapped reads (mapping quality >= 25) were used for further analysis (average deduplicated 23 millions of reads per sample). Peak locations were determined using the MACS algorithm (v2.1.0) with a cutoff of p-value = 1e-7. Peaks that were on the ENCODE blacklist of known false ChIP-Seq peaks were removed. Differentially modified locus of H3K27ac locus were identified by DiffBind package (v2.16). Peak annotation was performed by Homer program (v4.11). Promoter region cis-element prediction was performed by universalmotif package (v1.6.3).

### RNA extraction, quantitative real time PCR (qRT-PCR) and western blotting analysis

Total RNA was extracted using the RNeasy kit (Qiagen) according to the manufacturer’s protocol. For cDNA synthesis, reverse transcription was performed using M-MLV reverse transcriptase (Thermo Fisher Scientific) via manufacturer instructions. qRT-PCR was performed using QuantaStudio7 Real-Time PCR System (Applied Biosystems). Taqman probes used in this study were as follows. FOXA1 (Hs04187555_m1), CD274 (Hs00204257_m1). Relative gene expression change was calculated by the deltadeltaCt method. 18S ribosomal RNA was used as an endogenous reference. Western blotting was performed as described previously^3^. Primary antibodies were used as follows. FOXA1 (1:500, ab23738, Abcam), PD-L1 (1:1000, ab213524, abcam), GAPDH (14C10) (1:1000, #2118, Cell signaling).

### RNA-Sequencing of human tumors and UMUC1 cells

Briefly, RNA was extracted using RNeasy mini kit (Qiagen; Valencia, CA) according to manufacturer’s instructions. After ribogreen quantification and quality control with an Agilent BioAnalyzer, 2ug of total RNA underwent polyA selection and Truseq library (TruSeq™ RNA Sample Prep Kit v2) preparation. Briefly, samples were fragmented for 2 minutes at 94C before undergoing first strand and second strand cDNA synthesis. Libraries were amplified with 10 cycles of PCR and size-selected for fragments between 400 and 550 bp with a Pippin prep instrument (Sage Science). Samples were barcoded and run on a Hiseq 2500 in a 100bp paired end run, using the TruSeq SBS Kit v3 (Illumina). Sequence reads were aligned and counted for each gene by RSEM algorithm (<u>26</u>) with the STAR alignment program. An average of 30 million paired reads were achieved per sample, reads normalization and gene expression were analyzed using DESeq2. Gene pathway analysis was performed by GSEA.

## Acknowledgments

Supported in part by RSG 17-233–01-TBE from the American Cancer Society (D.J.D.), The W.W. Smith Charitable Trust (D.J.D.), the Ruth Heisey Cagnoli Endowment in Urology at Penn State College of Medicine, the Bladder Cancer Support Group at Penn State Health, the Leo and Anne Albert Institute for Bladder Cancer Care & Research, the Sloan Kettering Institute for Cancer Research Cancer Center Support Grant (P30CA008748), the SPORE in Bladder Cancer (P50CA221745) and R01CA233899 (H.A.).

## Notes

### Competing Interest Statement

The authors have declared no competing interest.

